# H3K4 methylation regulates development, DNA repair, and virulence in Mucorales

**DOI:** 10.1101/2023.06.05.543666

**Authors:** Macario Osorio-Concepción, Carlos Lax, Damaris Lorenzo-Gutiérrez, José Tomás Cánovas-Márquez, Ghizlane Tahiri, Eusebio Navarro, Ulrike Binder, Francisco Esteban Nicolás, Victoriano Garre

## Abstract

Mucorales are basal fungi that opportunistically cause a fatal infection known as mucormycosis (black fungus disease), which poses a significant threat to human health due to its high mortality rate and its recent association with SARS-CoV-2 infections. On the other hand, histone methylation is a regulatory mechanism with pleiotropic effects, including the virulence of several pathogenic organisms. However, the role of epigenetic changes at the histone level never has been studied in Mucorales. Here, we dissected the functional role of Set1, a histone methyltransferase that catalyzes the methylation of H3K4, which is associated with the activation of gene transcription and virulence. A comparative analysis of the *Mucor lusitanicus* genome (previously known as *Mucor circinelloides* f. *lusitanicus*) identified only one homolog of Set1 from *Candida albicans* and *Saccharomyces cerevisiae* that contains the typical SET domain. Knockout strains in the gene *set1* lacked H3K4 monomethylation, dimethylation, and trimethylation enzymatic activities. These strains also showed a significant reduction in vegetative growth and sporulation. Additionally, *set1* null strains were more sensitive to SDS, EMS, and UV light, indicating severe impairment in the repair process of the cell wall and DNA lesions and a correlation between Set1 and these processes. During pathogen-host interactions, strains lacking the *set1* gene exhibited shortened polar growth within the phagosome and attenuated virulence both *in vitro* and *in vivo*. Our findings suggest that the histone methyltransferase Set1 coordinates several cell processes related to the pathogenesis of *M. lusitanicus* and may be an important target for future therapeutic strategies against mucormycosis.

**Author Summary:** The knowledge regarding the role of epigenetic modification in regulating gene expression in early diverging fungi is scarce, despite they represent an important fraction of the fungal kingdom. The order Mucorales, which causes the lethal infection known as mucormycosis, is not an exception. There is an urgent need to enhance our understanding of the biology of these fungi to develop effective treatments for mucormycosis, which are currently absent due to the natural resistance of Mucorales to most antifungal drugs. This work represents the first investigation into the role of the methylation of lysine 4 on histone 3 (H3K4) in a mucoralean fungus. This was accomplished by the generation of deletion mutants in the *set1* gene, which encodes the specific H3K4 methyltransferase. Phenotypic analyses of these mutants suggest that H3K4 methylation regulates physiology, development, cell wall integrity, and DNA repair. Furthermore, our findings indicate that it also contributes to the virulence of *M. lusitanicus*, as strains lacking the *set1* gene exhibited shortened polar growth within the phagosome and attenuated virulence both *in vitro* and *in vivo*.

## Introduction

Mucorales are early diverging fungi that are considered saprophytic and globally distributed [1]. However, certain species can evolve as pathogens by adapting to different conditions, significantly compromising human health. Mucormycosis is an invasive infection caused by this group of fungi, particularly species of the genera *Rhizopus, Mucor,* and *Lichtheimia*, which infect host tissues and can spread to other organs [2–5]. Although it has traditionally been considered a disease exclusive to individuals with compromised immune systems resulting from cancer treatments, hematological disorders, and uncontrolled diabetes, it has also been detected in healthy individuals [6–8]. Mucormycosis can be acquired through inhalation, ingestion, or traumatic inoculation of Mucorales spores and can manifest in various clinical forms such as pulmonary, cutaneous, gastrointestinal, and rhinocerebral [9–11]. Despite the development of new antifungal agents, the incidence of mucormycosis has significantly increased in recent years, with alarming mortality rates of up to 90% [8,11–14]. In addition, most Mucorales species exhibit innate resistance and have evolved various multi-resistance pathways to commonly used therapeutic options, making mucormycosis a deadly disease [15–17].

*M. lusitanicus* is a suitable model organism for studying the molecular mechanisms underlying the physiology, development, pathogenesis, and virulence of Mucorales. The development of new molecular techniques and omics technologies has substantially contributed to identifying several pathways related to the virulence and antifungal resistance of Mucorales [17–19]. Furthermore, these mechanisms have facilitated a better understanding of the genetics and pathogenesis of Mucorales. In this context, the regulatory mechanisms based on histone post-transcriptional modifications have been scarcely studied in Mucorales [20], raising the consideration of epigenetic modifications as a new avenue to unveil the particularities of the mucoralean biology.

Epigenetic mechanisms modify gene expression transiently by finely modulating the genomic structure without altering the DNA sequence. These dynamic mechanisms are crucial in regulating the cellular response to different extra- and intracellular signals [21,22]. Major epigenetic modifications include post-translational modifications of histones, chromatin remodeling, RNA interference (RNAi), and DNA methylation [21–23]. The amino-terminal ends of histones serve as targets for numerous covalent modifications, such as methylation, acetylation, phosphorylation, and ubiquitination which finely tune chromatin status and gene expression [24–28]. Histone methylation is a chemical modification that occurs at specific lysines and arginines of the different histones; among the most studied are those of lysines 4, 9, 27, and 36 on histone H3, as marks for binding reader proteins that modulate gene transcription [29–33].

The biological role of histone methylation marks depends exclusively on the site and extent of methylation [34,35]. Histone 3 lysine 4 (H3K4) methylation plays an important role in the transcriptional activation of genes related to development, pathogenicity factors, secondary metabolism, and DNA repair in several pathogenic fungi [36–41]. H3K4 methylation is executed by the Set1 enzyme, a methyltransferase with a conserved SET domain initially identified in yeast [42–44]. In yeast, Set1 acts as the catalytic component of the Complex Protein Associated with Set1 (COMPASS) for the different extents of H3K4 methylation [35,45,46].

Mutations in *Set1* in *Saccharomyces cerevisiae* suppress H3K4 methylation, repressing transcriptional activity, which results in growth abnormalities and a deficiency in the DNA repair [28,47–51]. Similarly, *Set1* disruption in plant pathogenic fungi, including several *Fusarium* species, *Magnaporte oryzae*, and *Aspergillus flavus*, has been shown to cause loss of the H3K4 mark. This loss affects gene transcription, secondary metabolism, response to several stress types, hyphal growth, conidiation, appressorium formation, and virulence [36,37,40,51,52].

This H3K4 epigenetic mark also plays a crucial role in the pathogenesis of the human fungal pathogen *Candida albicans*. Deletion of *Set1* in this fungus results in total loss of H3K4 methylation, triggering hyperfilamentous growth in particular *in vitro* conditions, alterations of the cell surface, and reduced adherence to epithelial cells. Furthermore, murine model infectious assays showed that *set1* mutants could not develop an infection, as high animal survival rates were observed, suggesting that *set1* is critical for *Candida* pathogenesis through H3K4 methylation [39].

The lack of information about the role of histone modifications in the biology of Mucorales and in their ability to cause infection prompted us to analysis the function of *set1* gene, which codes for a Set1-type methyltransferase in *M. lusitanicus*. The deletion of *set1* confirmed the H3K4 methylation is strictly dependent on Set1 and revealed its crucial role in regulating development, DNA repair, and stress responses in *M. lusitanicus*. Furthermore, set1 is required for full virulence and proper polar growth after phagocytosis, as well as induced cell death of macrophages. These findings provide the first evidence of epigenetic modifications, specifically histone methylation, regulating the physiology and pathogenesis of *M. lusitanicus*.

## Results

### The genome of *M. lusitanicus* encodes Set1, an enzyme with histone methyltransferase activity

To identify the genes involved in H3K4 methylation, we inspected the proteomes of 20 representative species of the major fungal phyla using the *S. cerevisiae* the COMPASS components as the query (**Fig 1A** and **S1 Dataset**). Overall, the components of the COMPASS are highly conserved across fungi, with the only exception of the Cps15/Shg1, a non-essential component of this complex found exclusively in *S. cerevisiae* [53], which is also absent in animals (**Fig 1A**). All the other components are found in most of the fungal phyla, apart from the Cps25/Sdc1 component that is absent in two distant phylogenetic fungal groups: Zoopagomycota and Basidiomycota. The presence of all the components of the COMPASS in Mucorales suggests the importance of H3K4 methylation in this group of fungi. To understand the function of H3K4 in Mucorales biology, we focused on the *set1* gene of *M. lusitanicus*, which encodes the key methyltransferase component of the COMPASS. The predicted amino acid sequence of the only one *set1* homolog (JGI ID 1544069) of *M. lusitanicus* contains a putative SET domain, characteristic of known Set1 methyltransferases (**Fig 1B**). Furthermore, phylogenetic analysis revealed a high degree of homology between the Set1 enzymes from *M. lusitanicus* and other fungal species, with the *M. lusitanicus* methyltransferase clustering closely with the Set1 proteins from *C. albicans* and *S. cerevisiae* (**Fig 1C** and **S1 Table**). Taken together, these findings suggest that Set1 likely has a conserved activity in methylating histone 3 lysines 4, which may regulate a variety of cellular processes in *M. lusitanicus*, similar to the function of characterized methyltransferases Set1 in other fungi [38,39]. To test this hypothesis, we deleted the *set1* gene in the strain MU402 strain (*pyrG*ˉ, *leuA*ˉ) using the strategy of replacement by double homologous recombination (**Fig 2A**) [54]. Endpoint PCR analysis of two candidate mutant strains, which derived from independent genetic transformations, showed that *pyrG* replaced the *set1* gene in homokaryosis. This was confirmed by the observation of only a fragment of 5.1 kb corresponding to the mutant locus (**Fig 2B**). These mutants were designated as *set1*-3Δ and *set1*-4Δ for further analysis (**S2 Table**).

**Fig 1.**
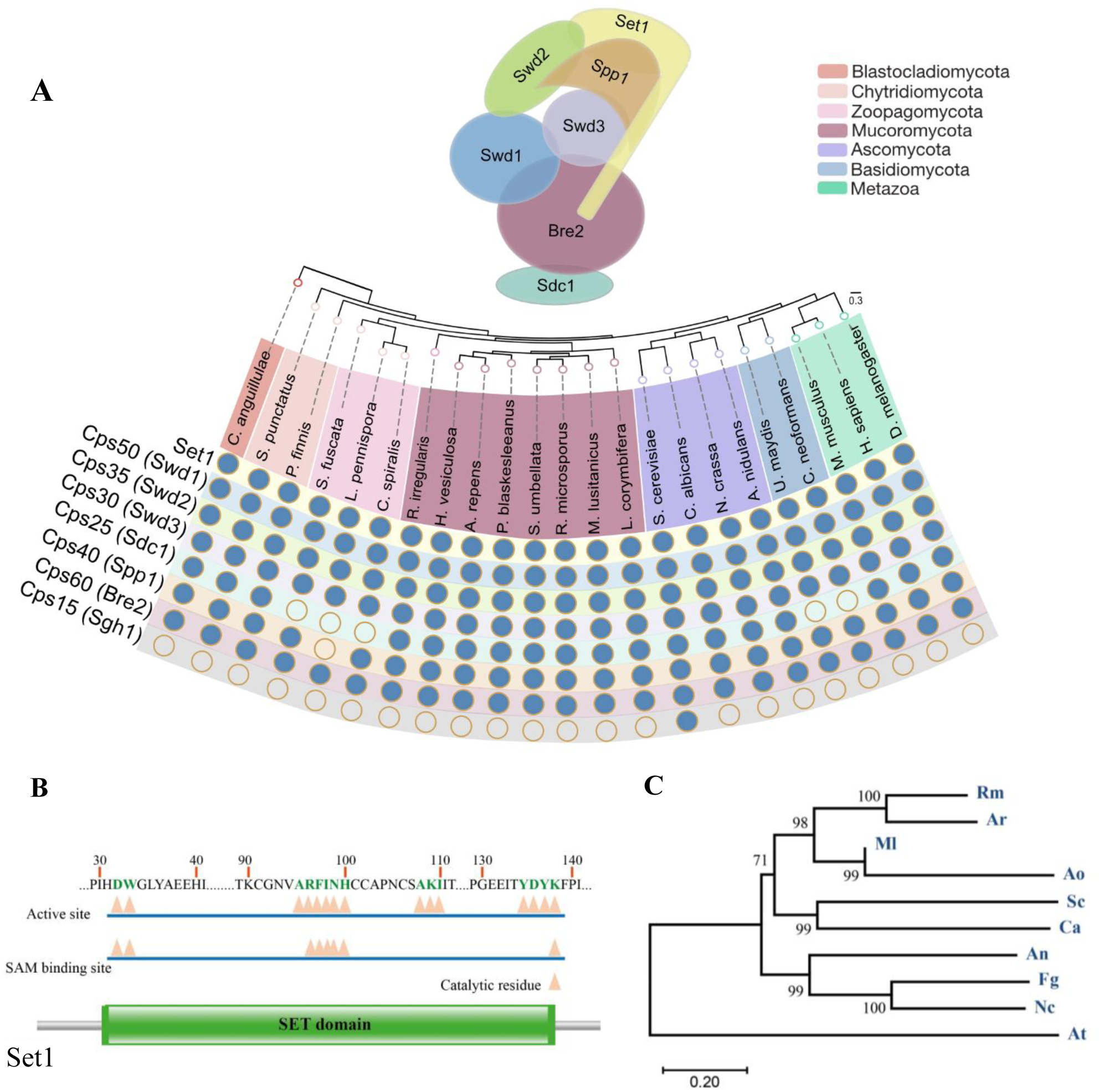
**Components of the COMPASS complex**. **A)** Conservation of the COMPASS complex in fungi and animals. **B)** Schematic representation of *M. lusitanicus* Set1 containing the putative SET domain, which includes catalytic residue, SAM binding site, and active site with the corresponding amino acids. **C)** Phylogenetic analysis of Set1 homologs (**S1 Table**) from *M. lusitanicus* (Ml) and other fungal species such as *Rhizopus microsporus* (Rm), *Aspergillus rouxii* (Ar), *Aspergillus ossiformis* (Ao), *S. cerevisiae* (Sc), *C. albicans* (Ca), *Aspergillus nidulans* (An), *Fusarium graminearum* (Fg) and *Neurospora crassa* (Nc) conducted with the MEGA version X software using the neighbour-joining method. *A. thaliana* (At) was included as an outgroup. The phylogenetic tree shows the bootstrap values in each node from 1000 replicates. The conserved SET domain was determined by Pfam (http://pfam.xfam.org/).

**Fig 2.**
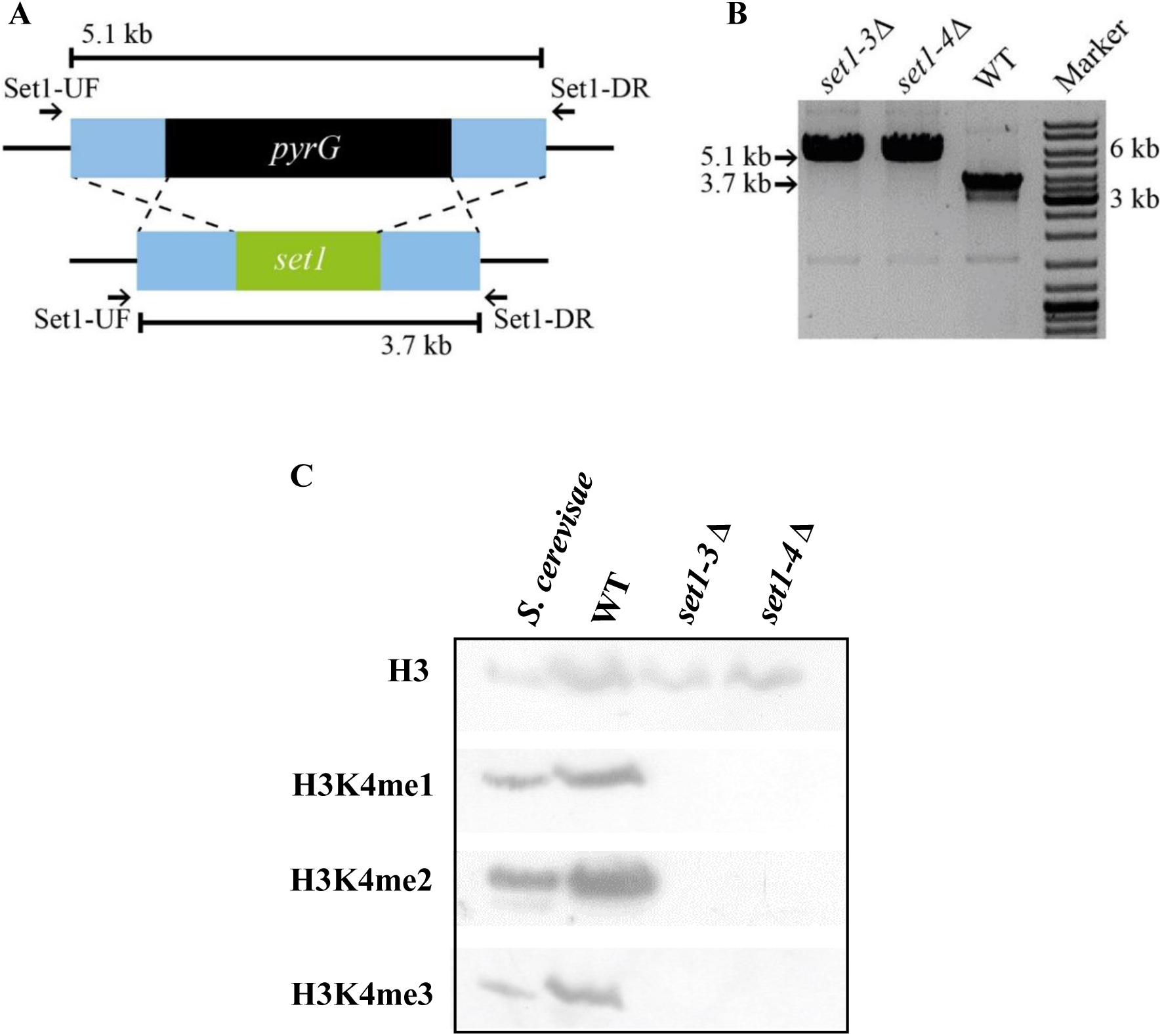
**Functional analysis of the gene *set1* of *M. lusitanicus*. A**) Scheme of mutant locus after *set1* locus replacement on the genome of wild-type strain by the *pyrG* construct by homologous recombination. The recombination sites on the genome are marked with dashed lines. The binding sites of oligonucleotides used to check genic deletion and the size of amplified fragments are denoted on the diagram. **B**) PCR products of *set1* locus of wild-type strain MU636 (WT) and selected *set1* mutants were amplified with the primers Set1-UF and Set1-DR (**S3 Table**) and separated by electrophoresis. The expected band (black arrows) for the mutant and wild-type locus corresponded to 5.1 kb and 3.1 kb, respectively. **C)** H3K4 methylation in the wild-type strain (wt) and *set1* mutants was determined by western blot analysis by using specific antibodies against H3K4 mono-(me1), di-(me2) and trimethylation (me3). As a control, *S. cerevisiae* protein extract was included. The protein samples were also incubated with anti-histone H3 antibody as a loading control.

To determine if Set1 is the methyltransferase responsible for H3K4 methylation, we used the western blotting approach and fungal mono-, di-, and trimethylated forms of H3K4-specific antibodies to determine H3K4 methylation levels in each mutant. In these western blottings, H3K4 monomethylation (me1), dimethylation (me2) and trimethylation (me3) were not detected in the two *set1*-independent mutants (**Fig 2C**). These results demonstrated the essential role of Set1 in H3K4 methylation in *M. lusitanicus*.

### *set1* is involved in growth, asexual sporulation, cell wall integrity and DNA-repair

To investigate the role of *set1* on the physiology and development of *M. lusitanicus*, we performed a phenotypic analysis of *set1* mutants. Deletion of the *set1* gene has a negative impact on the growth of *M. lusitanicus*, as both *set1* mutants displayed a smaller colony diameter compared with the wild-type strain MU636, measured at 24-h intervals up to 72 h (**Figs 3A and 3B**). Moreover, the lack of *set1* reduced the production of asexual spores in both mutants, generating less than half the spores/cm^2^ compared to the WT (p=0,005) (**Fig 3C**). These mutants also showed increased sensitivity to cell wall stress induced by the presence of sodium dodecyl sulfate (SDS). We dropped appropriate spore concentrations (10^4^, 10^3^, 10^2^, and 10 spores) of each strain on YNB plates containing 0.002% SDS. The ability of the strains to form colonies in the presence of SDS was used to determine their sensitivity levels. Our results showed that both *set1* mutants were highly sensitive to SDS compared to the wild-type strain, with complete inhibition of growth at low spore concentrations (10^3^, 10^2^, and 10 spores) (**Fig 4A**). This phenotype was further supported by the lower percentage of *set1* mutant spores that form colonies in medium seeded in plates with SDS (**Fig 4B** **and S1 Fig**). These results suggest that the activity methyltransferase of Set1 is important for appropriate mycelial growth, sporulation process, and integrity of the cell wall of *M. lusitanicus*.

**Fig 3.**
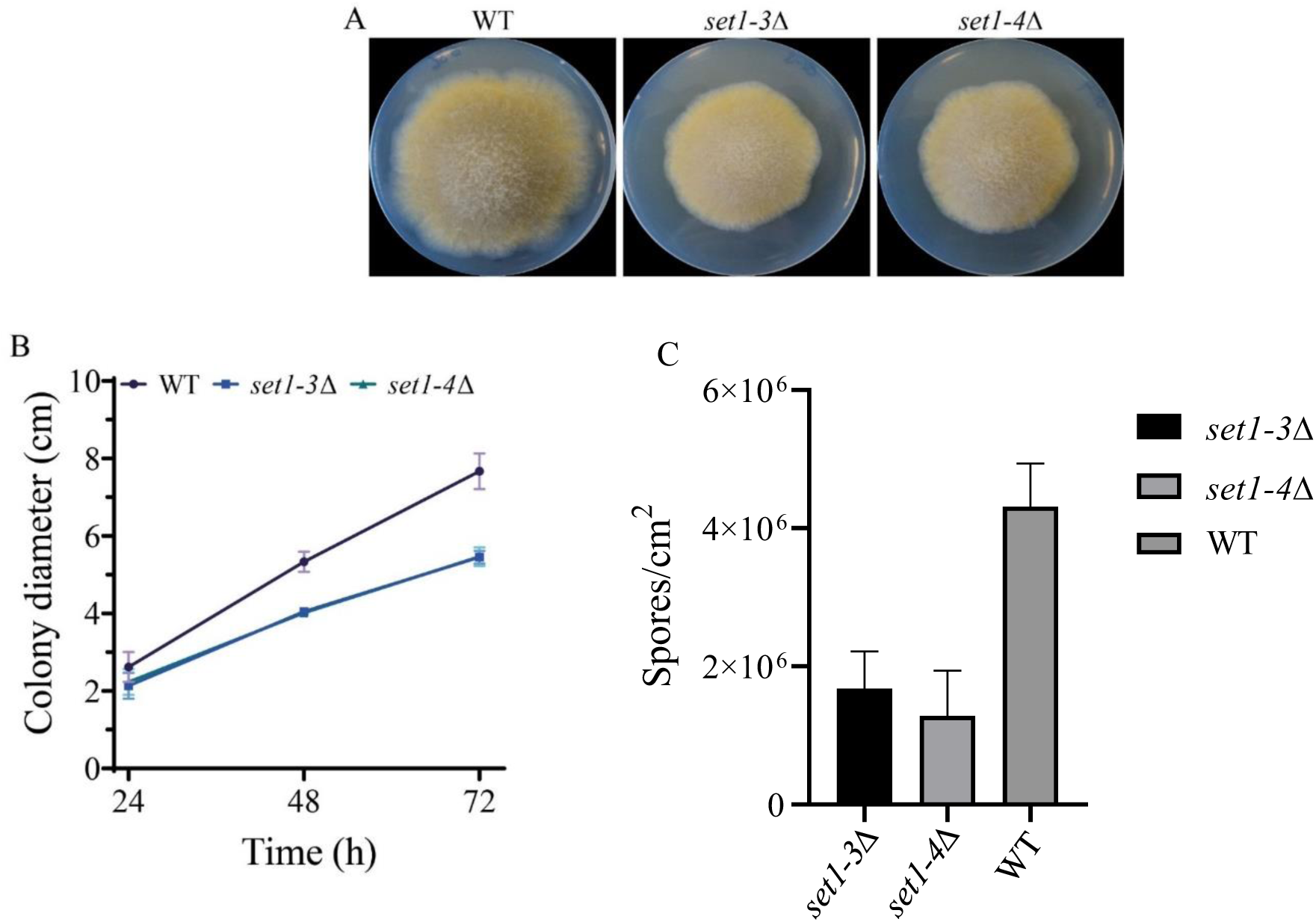
**The deletion of *set1* alters colony growth of *M. lusitanicus***. **A**) Fungal colonies of the wild-type strain MU636 (WT) and *set1* deletion mutants (*set1-3*Δ and *set1-4*Δ) grown on YPG medium plates pH 4.5 at 26°C for 3 days under constant illumination. **B**) Graph displays the radial growth of the WT and *set1Δ* strains measured every 24 h for 72 h. The line bars in each time point of the growth kinetic represent the standard error from the three biological experiments. **C**) Production of vegetative spores in *set1* mutants and WT on solid YPG pH 4.5.

**Fig 4.**
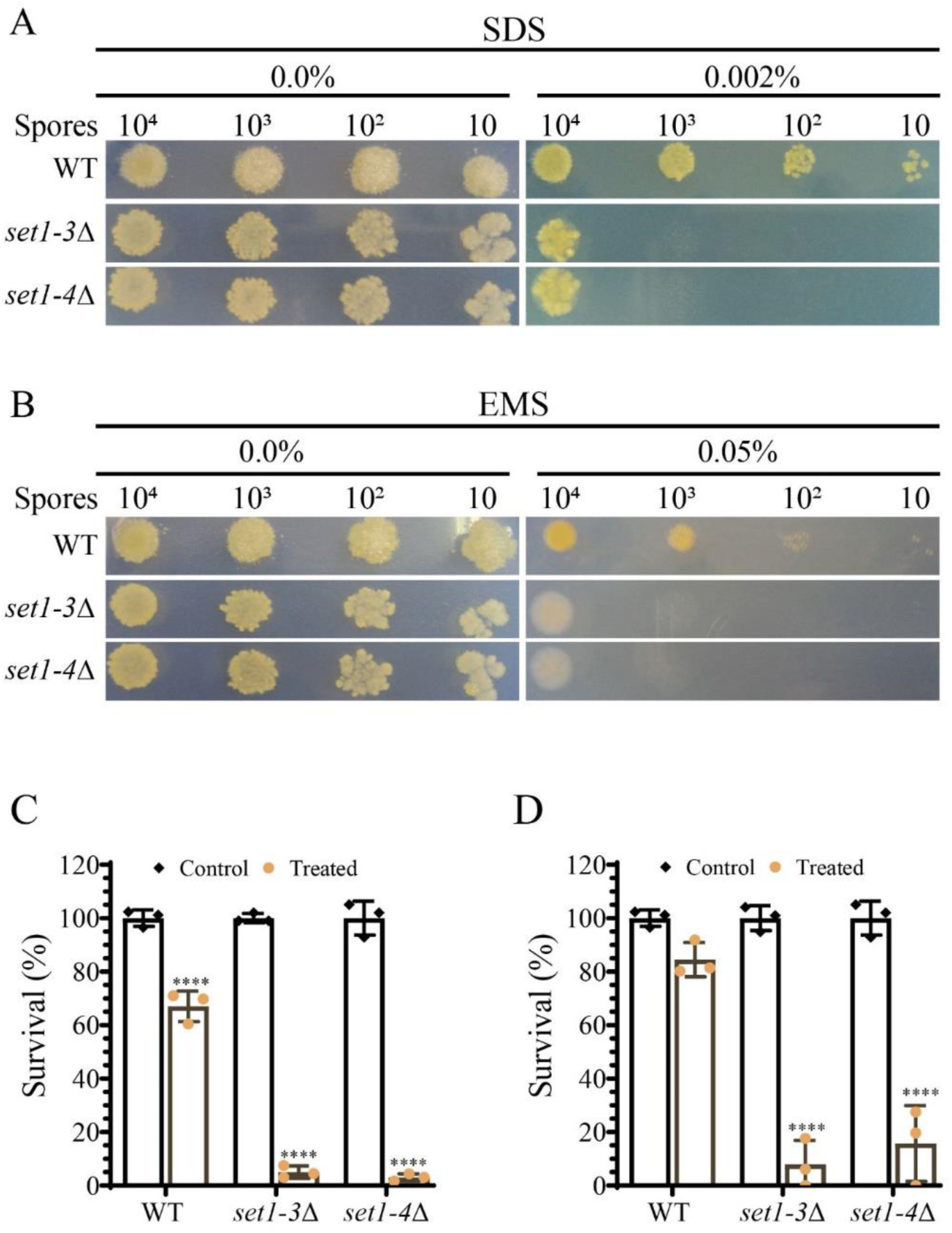
***set1* knockout strains are sensitive to SDS and EMS**. **A** and **B**) Growth of wild-type strain MU636 (WT) and *set1Δ* strains from different concentrations of spores spotted on YNB medium, YNB medium amended with SDS (0.002%) or EMS (0.05%). **C** and **D**) Survival rate of *M. lusitanicus* strains after treatment with SDS (0.002%), EMS (0.05%), and control conditions. The cultures were placed at 26 °C by 48 hours. Data were analyzed by two-way ANOVA and asterisks above the charts indicate significant differences (**** P < 0.0001).

Since Set1 has been involved in DNA repair in *S. cerevisiae* [55], we evaluated the involvement of *M. lusitanicus set1* in DNA repair by analyzing the sensitivity of *set1*Δ strains to the alkylating agent ethyl methanesulfonate (EMS) and UV light, which induces dimerization of DNA bases [56,57]. Spores of the *set1*Δ strains either grown in the presence of 0.05% EMS or exposed to a UV pulse (10 mJ/cm2) developed fewer colonies and exhibited drastically reduced survival levels compared to the wild-type strain (**Figs 4C-4F and** **S1 Fig**), suggesting that H3K4 methylation plays an important role in DNA damage repair in *M. lusitanicus*.

### The lack of *set1* reduces virulence of *M. lusitanicu*s

Set1 enzymes have been involved in the virulence of a few fungal pathogens belonging to the Ascomycota phylum [36,37,39,40,51,52], but its role has not been analyzed in other fungal groups. Therefore, we assessed the role of *set1* in the pathogenesis of *M. lusitanicus* using the *Galleria mellonella* infection model [58]. Healthy larvae of *G. mellonella* were injected with spores of the wild-type strain or the *set1* mutant strains and their survival was monitored daily for 6 days. Intriguingly, both *set1* deletion mutants exhibited reduced virulence compared to the wild-type strain (p = 0.0348 and p = 0.0160, respectively) (**Fig 5**), suggesting that the Set1 enzyme plays a critical role in the pathogenicity of *M. lusitanicus*.

**Fig 5.**
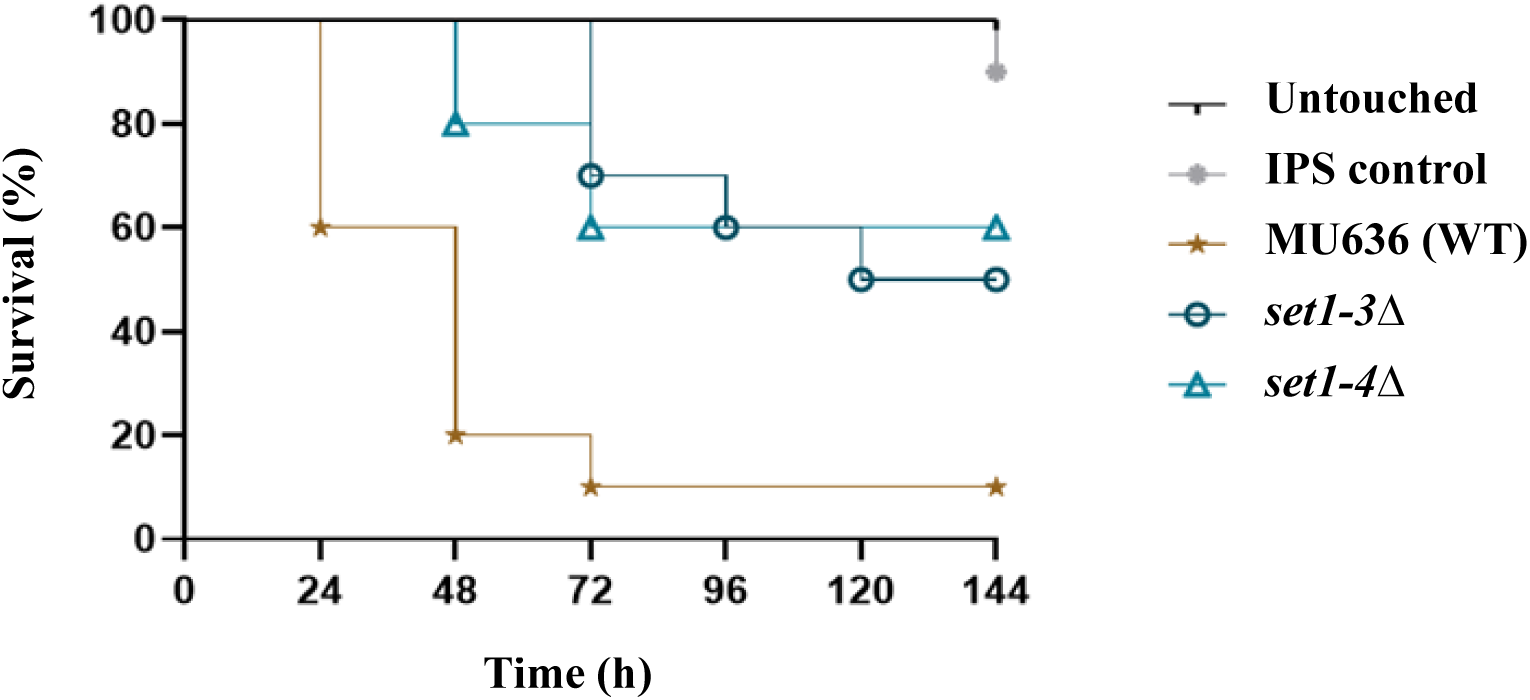
Set1 is essential for *M. lusitanicus* virulence. Virulence of *M. lusitanicus* strains in *G. mellonella* model host inoculated with 10^6^ spores of the WT strain (MU636) and *set1* mutants through the last pro-leg into the hemocoel. Percentage survival values were plotted for 6 days. Untouched larvae and larvae injected with IPS were used as control. Survival curves were statistically analyzed by log-rank (Mantel-Cox) test, utilizing GraphPad Prism 8.0.2 software. The difference in survival between *set1* mutants and wild-type strain was statistically significant (P values ≤ 0.05).

To investigate the contribution of Set1 protein to the ability of *M. lusitanicus* to survive phagocytosis during interaction with macrophages, we challenged spores of *set1-3*Δ and *set1-4*Δ mutants with mouse macrophages for 5.5 h of co-culture and subsequently plated them on MMC agar plates. Similar to the WT strain, both *set1* mutants retained the ability to counteract the cytotoxic environment during phagocytosis, as evidenced by the lack of damage in the grown colonies (**S2A Fig**). This finding suggests that the regulatory mechanism governing *M. lusitanicus* survival during macrophage phagocytosis does not depend on the Set1 enzyme. However, we observed that phagocytized spores from *set1* deletion strains developed a shorter germ tube after 5.5 h of interaction with macrophages (**S2B Fig**). In contrast to the wild-type strain, the polarity rate, calculated from the length of the emerged hyphae and spore width, was significantly lower in the *set1-3*Δ and *set1-4*Δ mutants in comparison to the wild-type strain (**Fig 6A**), indicating that Set1 is required for appropriate germ tube development during phagocytosis.

**Fig 6.**
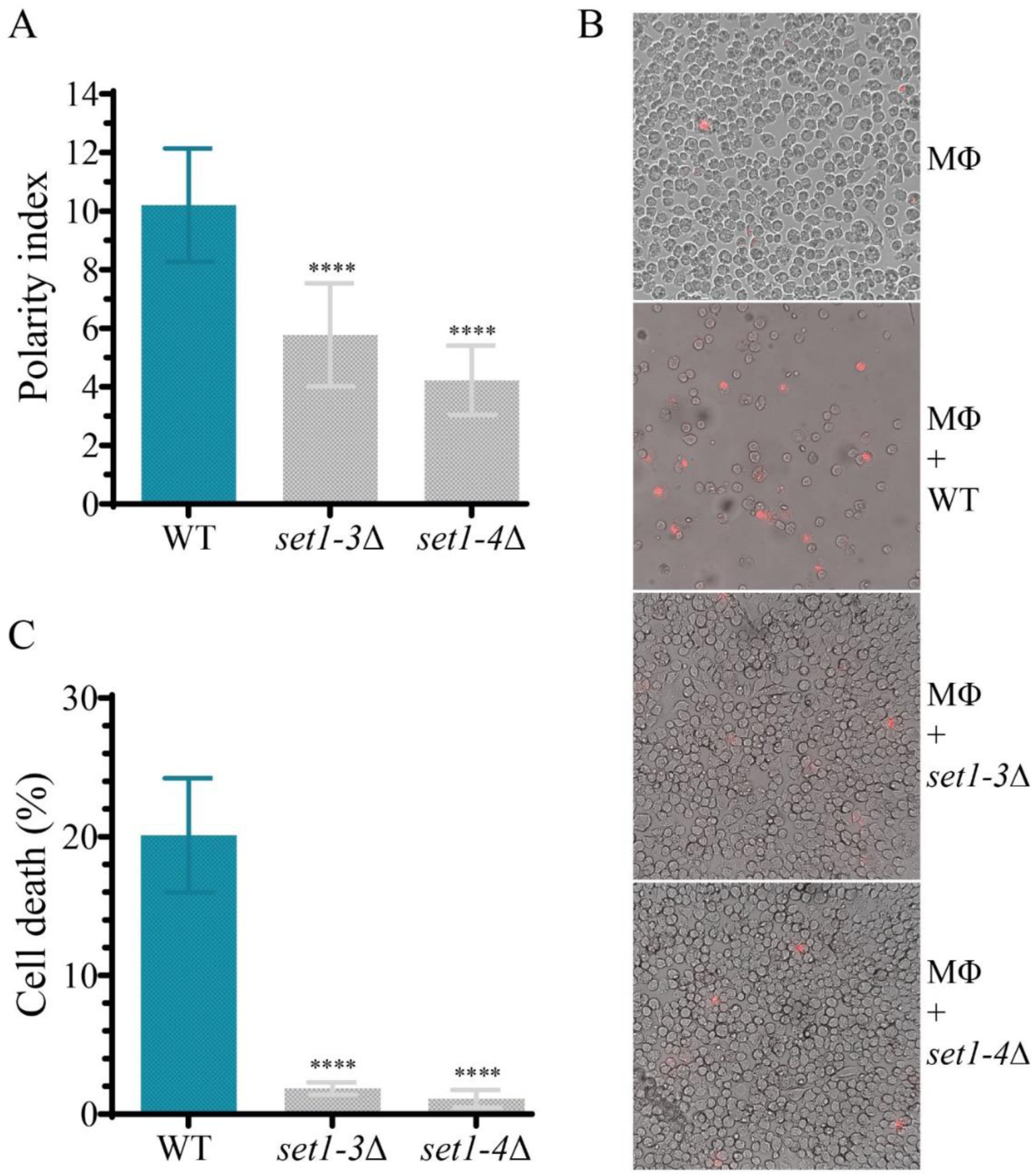
***set1* disruption results in a reduction of polarity index and ability to induce macrophage death of *M. lusitanicus***. **A**) Polarity index of the wild-type MU636 (WT) and *set1* deletion strains determined from phagocytosed spores by mouse macrophages (J774A.1) after 5.5 hours of incubation. **B**) Images of live-cell microscopy of mouse macrophages (Φ) co-cultured for 24 h with spores of WT and *set1Δ* strains. The dead macrophages were stained with propidium iodide (PI). Macrophage non-interacting with spores served as a control. **C**) Macrophages cell death at 24 hours of interaction with *M. lusitanicus* spores estimated by determining percentage of PI-stained macrophages in the fluorescent images. The charts display means ± SD based on three biological repetitions. The statistical differences were obtained by one-way ANOVA and indicated by asterisks (****P < 0.0001).

The decreased polar growth during interaction with macrophages could potentially impact *M. lusitanicus* pathogenesis. To test the role of Set1 in macrophage cell death, we performed experiments on the interaction between *set1* mutant spores and mouse macrophages. Cell death was determined 24 h later by adding propidium iodide (PI), which stains dead cells or those with a damaged cell membrane. We observed fewer dead macrophage cells stained with PI in cultures with *set1-3*Δ and *set1-4*Δ mutants compared to the wild-type strain (**Fig 6B**). Furthermore, the percentage of macrophage cell death, calculated from PI-positive cells and the total number of cells, was remarkably lower in the strains lacking the *set1* gene compared to the WT strain (**Fig 6C**). These results indicated that *M. lusitanicus* mediated killing of macrophages is regulated by the Set1 protein, likely through the methylation of H3K4.

### Genes regulated by *set1*

We performed a transcriptomic analysis to explore the effect of the lack of H3K4 methylation on the gene expression profile of *M. lusitanicus*. Messenger RNA was isolated and deep sequenced from the wild-type control and *set1* mutant strains after 24 h of growth on solid rich (YPG) medium. Transcriptomic analysis revealed a limited effect of H3K4 methylation on gene expression profiles in these growth conditions (**Fig 7A**, **S2 Dataset**). A total of 403 genes were differentially expressed (DEGs) in the *set1* mutant strain: 343 up-regulated genes and 60 down-regulated genes. This was a surprising result as H3K4 methylation is considered an epigenetic mark related to actively expressed genes in fungi [36–41] and suggests that in *M. lusitanicus,* this regulation may affect other transduction pathways regulators that control the expression of the up-regulated genes. Unfortunately, more than half of the genes (243/403) lacked EuKaryotic Orthologous Group (KOG) annotation, which hampered our understanding of their contribution to the phenotypes observed. The annotated genes were mainly related to metabolism (94/160), highlighting the presence among top down-regulated DEGs of a putative phosphoinositide phosphatase (ID1537005), that showed a 41% of identity with Sac1 of *S. cerevisiae* (ScSac1). The blast analysis of this putative *M. lusitanicus* Sac1 (MlSac1) revealed that both were the best reciprocal hits in *S. cerevisiae* proteome, and the superimposition of MlSac1 predicted structure with ScSac1 suggested that they were true homologs (**Fig 7B**). Sac1 protein plays an essential role in the vesicle trafficking of *S. cerevisiae* being involved in protein secretion and cell wall maintenance [59]. This phosphoinositide phosphatase modulates phosphatidylinositol 4-phosphate concentration gradients driving the membrane homeostasis [60]. The down-regulation of MlSac1 due to *set1* mutation could unbalance the secretory pathways *of M. lusitanicus,* affecting the exportation of virulence factors and the proper composition of the cell membrane and cell wall.

**Fig 7.**
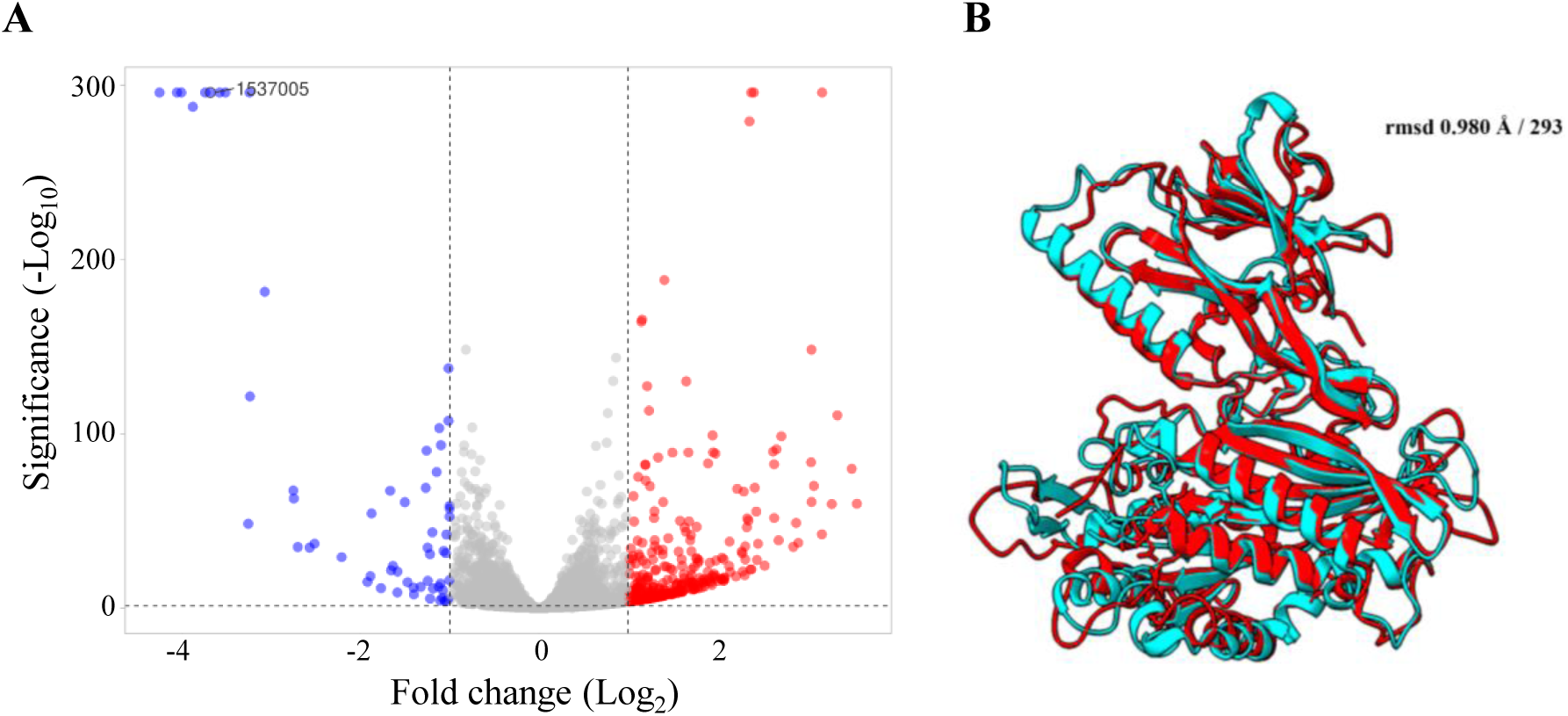
**Genes regulated by *set1***. **A**) Volcano plot of differentially expressed genes (DEGs) identified between a *set1* mutant and a wild-type strain. The red dots denote up-regulated genes, the blue dots denote down-regulated genes, and the gray dots denote the genes with expression changes or significance below threshold (See material and methods). **B**) Superimposition of ScSac1 (PDB <u>3LWT</u>) in red and the predicted structure for MlSac1 in blue. The rmsd and the number of aligned residues are indicated.

## Discussion

Identifying novel regulators involved in the pathogenesis of Mucorales is crucial and could help to develop specific antifungal treatments for early mucormycosis therapy. Set1 is a histone methyltransferase of the COMPASS protein complex that catalyzes the addition of methyl groups to lysine 4 on histone 3 via its SET domain [35,53]. Generally, H3K4 methylation has been associated with the transcriptional activation of genes involved in various biological events in eukaryotes [37,48,61–63]. In several fungal species, the Set1 protein is required for cellular responses to stressful environments and pathogenesis [36–39]. However, the role of the methyltransferase Set1 in Mucorales has not been investigated. In this study, we characterized the function of the unique *set1* gene in *M. lusitanicus,* providing new insights into the regulatory mechanisms of morphogenesis and pathogenesis in this fungus at the epigenetic level. The *set1* gene of *M. lusitanicus* encodes a putative histone methyltransferase that shares high similarity to well-characterized Set1 of *S. cerevisiae* and *C. albicans*. Set1 deletion in *C. albicans* causes growth defects, reduces adherence to epithelial cells, and suppresses histone 3 lysine 4 (H3K4) methylation, leading to attenuated virulence [39]. Similarly, in *S. cerevisiae*, disruption of *SET1* results in a loss of H3K4 methylation and a reduction in gene transcription [48]. Several phytopathogenic fungi have reported similar findings [36–38,40]. The amino acid sequence of *M. lusitanicus* Set1 harbors the SET domain characteristic of known histone methyltransferases Set1, which is essential for H3K4 methylation and supports several biological events observed in various eukaryotic organisms. As an indication of transcriptional activation, H3K4 methylation is a conserved epigenetic modification found from yeast to humans. It is widely believed that H3K4 methylation is catalyzed by the methyltransferase Set1/COMPASS [64]. In the phytopathogenic fungus *Fusarium graminearum*, FgSet1 is responsible for H3K4 mono-, di-, and trimethylation [36], whereas Set1 of *Aspergillus flavus* (AflSet1) only produce di-and trimethylation [52]. To investigate the biological role of Set1 in histone methylation in the fungus *M. lusitanicus*, western blotting analysis was performed in this study. The results of the immunoblotting analysis showed that similar to FgSet1, Set1 primarily regulates the mono-, di-, and trimethylation of H3K4. In contrast to AflSet1, however, Set1 is also responsible for H3K4 monomethylation in *M. lusitanicus*. These findings indicate that the methylation activity of Set1 is relatively conserved across different fungal species but shows functional differences related to the number of methylations.

In *M. lusitanicus*, as in other filamentous fungi and yeast, Set1 is associated with vegetative growth. The absence of Set1 resulted in a significant decrease in colony growth, indicating that Set1 is involved in this process. Moreover, our data suggest that the pathways regulating sporulation are also linked to Set1, as mutants and wild-type strains showed differences in spore production. In several phytopathogenic fungi species, Set1 is essential for spore formation [41], similar to the case in *M. lusitanicus*, indicating functional conservation among different species within the fungal kingdom.

Little information exists on the interplay between Set1, H3K4 methylation, and stress signal response. Our genetic replacement experiments demonstrated that the *set1* gene is required for *M. lusitanicus* to respond to cell wall-disrupting agents. The *set1* deletion mutants were highly sensitive to SDS, indicating that the cell wall stress signaling pathway is compromised in the absence of Set1, which ultimately compromises cell survival (**Figs 2A and 2C**). In *Fusarium verticillioides*, Set1 mediates the phosphorylation of the mitogen-activated protein kinase Mpk1 and Hog1 orthologs for preserving the cell wall integrity and responding to stress signals [40]. Therefore, it is plausible that the deregulation of this process in *M. lusitanicus* results from the lack of the *set1* gene.

Moreover, the repair of distinct types of DNA lesions was also impaired in *M. lusitanicus*. EMS reagent generates alkylated bases, which are removed from DNA by base excision [65]. The *set1* mutant strains were more susceptible to EMS, supporting the notion that the repair mechanism of damaged nucleotides is deficient in these mutants due to the deletion of the *set1* gene. Interestingly, these mutants also lose the ability to survive the genotoxicity of UV irradiation, as observed in our experiments. Applying UV light induces the formation of thymine dimers in DNA, and inaccurate repair by base excision leads to cell death [66]. These findings suggest that the Set1 protein plays an important role in different repair pathways, including cell wall and DNA lesion repair, and may regulate the expression of genes related to cell damage repair through its methyltransferase activity.

In contrast to the sensitivity of the Set1-deficient strains to chemical compounds that induce cell damage, these strains do not exhibit impairment in their response to survive phagocytosis during interaction with mouse macrophage cells. Within the phagosome, fungal spores are subjected to a stressful environment, including nutrient deprivation, antimicrobial peptides, acidification, and oxidative stress, which are known to affect their survival [67,68]. Interestingly, the colonies of the *set1* mutants presented healthy development after being challenged with macrophages, indicating that distinct defense strategies operate correctly to resist the macrophage attack. Previous studies have demonstrated the ability of some Mucorales species to germinate inside macrophages as a survival process and induction of cell death [69,70]. In this sense, the *set1* knockout strains displayed late germination supported by lower values of polarity index during the macrophage phagocytosis, suggesting an important role of the Set1 enzyme in the timely development of germ tube inside the phagosome. The *set1* removal probably leads to a breakdown in the regulatory pathways of germination of *M. lusitanicus* in response to macrophages *in vitro*. *M. lusitanicus* strains impaired in germination or with shortened polar growth are known to be less virulent [68]. After analyzing the virulence *in vitro* of *set1* mutants, we found that they present attenuated virulence due to the lower number of PI-stained macrophages observed in the fluorescent images, indicating a minor percentage of cell death. Thus, although the mutants can resist phagocytosis, they cannot kill macrophages at the same level as the wild-type strain. The attenuated virulence *in vitro* of these mutants was supported by infection experiments *in vivo* in *G. mellonella*, where lower mortality rates were observed in strains lacking the *set1* gene. These findings highlight the contribution of Set1 to the pathogenesis and virulence of *M. lusitanicus*. Previous studies have demonstrated that this fungus activates cell death by activating specific apoptosis-related genes in macrophages [68]. Therefore, the deregulation of this process could occur in the *set1* mutants and partially explain the reduced virulence. However, additional studies are required to demonstrate this hypothesis. Similar to our results, Set1 plays an essential role in virulence in several species of phytopathogenic fungi, and strains lacking the *set1* gene lose the ability to infect maize plants and defects in the toxin biosynthesis [40].

Transcriptomic analysis of *set1* mutants revealed the genes repressed and activated by this gene (**Fig 7**). The most striking result of this analysis was the larger number of genes that were repressed compared to those that were activated, which contrasts with the general activation of gene expression seen in other fungi. There are two potential explanations for these findings. Firstly, Set1 primarily activates gene expression, as seen in other organisms, but many activated genes act as repressors of different gene pathways. Alternatively, the less likely hypothesis is that the methylation of H3K4 by Set1 has a dual function, as observed with H3K9 methylation. Previous studies showed that the trimethylation of H3K9 is associated with transcriptional repressed loci, whereas dimethylation is associated with transcriptional activation [36]. Another interesting result from the transcriptomic analysis was the high number of genes regulated by Set1 that are involved in metabolism. This finding could be linked to the observed decrease in virulence both *in vitro* and *in vivo*. A metabolic outfit affected by the lack of Set1 could result in a delay in germ tube growth. This delay, in turn, would represent a disadvantage for the fungus during interaction with phagocytic cells, which could explain the overall decrease in virulence observed in *set1* mutant strains.

## Material and Methods

### Fungal strains and growth conditions

*M. lusitanicus* strain MU402 (*pyrGˉ*, *leuAˉ*) [71] was employed as a recipient strain during the process of genetic transformation to generate the *set*1 mutants (**S2 Table**). The strain MU636 (*leuAˉ*) [72], derived from MU402, was used as a wild-type strain for the different experiments throughout this research.

All the *M. lusitanicus* strains were grown at 26°C on yeast peptone glucose agar plates (YPG; 3 g/L yeast extract, 10 g/L peptone, 20 g/L glucose, 15 g/l agar), pH 4.5, under illumination conditions for sporulation and colony growth measurement. The colonies grown after protoplast transformation and the spores recovered from macrophages lysates were plated on Minimal Medium with Casaminoacids (MMC; 10 g/L casaminoacids, 0.5 g/L yeast nitrogen base without amino acids and ammonium sulfate, 20 g/L glucose, 15 g/L agar), adjusted to pH 3.2, and supplemented with niacin (1 mg/ml) and thiamine (1 mg/ml) to isolate homocaryotic strains and evaluate growth fitness, respectively. *M. lusitanicus* cultures grown on Yeast Nitrogen Base (YNB; 1.5 g/L ammonium sulfate, 1.5 g/L glutamic acid, 0.5 g/L yeast nitrogen base without amino acids and ammonium sulfate, 10 g/L glucose, and 15 g/L agar) with pH 3.0 at 26°C, supplemented with niacin (1 mg/ml), thiamine (1 mg/ml), and leucine (20 mg/L) were performed to examine the effect of ethyl methanesulfonate (EMS), sodium dodecyl sulfate (SDS), ultraviolet light (UV), and hydrogen peroxide (H2O2). Cell cultures for the experiments on macrophage cell death and polarity index were performed in Leivobitz L-15 medium (Biowest, Minneapolis, MN, USA) at 37°C, with 10% fetal bovine serum (FBS) and 1% penicillin/streptomycin (Gibco). Propidium iodide ReadyProbes reagent (Invitrogen) was used to stain macrophages and estimate cell death (2 drops per 10^6^ cells/ml).

### Phylogenetic analysis and ortholog search

Proteomes of 23 representative species were retrieved from the Joint Genome Institute (JGI) Mycocosm genome portal [73] and Uniprot [74]. Sequences of *S. cerevisiae* Set1, Cps15 (Sgh1), Cps25 (Sdc1), Cps30 (Swd3), Cps35 (Swd2), Cps40 (Spp1), Cps50 (Swd1) and Cps60 (Bre2) proteins were queried against the selected proteomes using iterative HMMER jackhmmer searches (E-value ≤ 1×10-3) (v3.3.2) (http://hmmer.org). A reciprocal BLASTp search (v2.10.1) [75] was conducted, and sequences that failed to produce a hit were discarded. An additional search using the Pfam-A database [76] using HMMER hmmscan (v3.3.2) (http://hmmer.org) served to remove hits that lacked the set domain (Set1 orthologs), WD/WD40 domains (Swd1, Swd2, and Swd3 orthologs), PHD-finger (Spp1 orthologs), Dpy-30 (Sdc1 orthologs), COMPASS Sgh1 (Sgh1 orthologs) and SPRY (Bre2 orthologs). A final list and a matrix including information about the presence or absence of the putative orthologs were generated (S1 Dataset). A Species tree was generated after analyzing the 23 proteomes with OrthoFinder (Inflation factor, -I 1.5) [77]. We identified 37040 protein orthogroups, of which 1200 had all species present. Sequences contained in single-copy orthogroups were aligned using MAFFT [78], and the phylogenetic species tree was obtained using RAxML [79] (PROTGAMMAWAGF substitution model) with 100 bootstrap replicates.

### Generation of *M. lusitanicus set1* knockout strains

#### Construction of recombinant fragment

To construct the recombinant fragment for *set1* gene disruption, 1 kb sequences of upstream and downstream regions from *set1* open read frame (ORF) and 2 kb sequence of *pyrG* selection marker were amplified by PCR using appropriate primers (**S3 Table**). Later, resulting PCR products were subjected to overlapping PCR employing specific oligonucleotides (**S3 Table**) to generate the construct, which consists of the *pyrG* selectable marker surrounded by 5′and 3’ ends of the *set1* gene. The PCR amplifications were performed with Herculase II fusión DNA Polymerase following supplier recommendation (Agilent, Santa Clara, CA, USA).

#### Genetic Transformation of *M. lusitanicus*

The protoplast preparation was conducted to genetically transform the recipient strain MU402 (*pyrGˉ*, *leuAˉ*) according to the previously established protocol [80]. The genetic transformation of the construct into strain MU402 (*pyrG*ˉ, *leuA*ˉ) was performed by the protoplasts electroporation to replace the *set1* locus by double homologous recombination, as has been previously reported [80]. Colonies developed from transformed protoplasts were transferred to a minimal medium with aminoacids (MMC), pH 3.2, supplemented with niacin (1mg/ml) and (1 mg/ml) by several growth cycles to select homokaryons. The integration of the construct into the target locus and homokaryosis was checked by amplifying the entire recombinant fragment by PCR using specific oligonucleotides (**S3 Table**).

### Protein extraction and detection

To prepare cell extracts from wt and *Δset1* strains, 100 mg of mycelium in 300 µl lysis buffer (8M Urea, 5% w/v SDS, 40mM Tris-ClH pH6.8, 0.1mM EDTA, 0.4 mg/ml bromophenol blue, 1% β-mercaptoethanol, 5mM PMSF, and 2% Protease Inhibitor Cocktail from Merck) were lysed with 0.5 mm Zirconia/Silica Beads in a FastPrep-24^TM^ homogenizer, centrifuged at 20000x g for 10 min at 4° C, and the supernatant was collected. 30 µl of histone extracts were resolved on 10% SDS-PAGE and transferred to a nitrocellulose membrane using the Semi-Dry Electroblotting Unit System (Sigma-Aldrich). Histone H3 and modified residues were detected with antibodies against H3 (Abcam, ab176842), H3K4me1 (EpigenTek, C10005-1-3K4M), H3K4me2 (Abcam, ab176878), or H3K4me3 (EpigenTek, C10005-1-3K4T).

### Phenotypic analysis of *set1* knockout mutants

To test the sporulation and colony growth of *set1* mutants (**S2 Table**), 500 fresh spores from each strain were inoculated in the center of solid YPG pH 4.5 and incubated at 26°C for 72 h in 12 h dark-light cycles. The number of produced spores was estimated using a Neubauer chamber and normalized with the growth area corresponding to each *M. lusitanicus* strain. For radial growth, the colony diameter from fungal strains was registered each 24 h by 3 days.

### Sensitivity testing

To determine the *M. lusitanicus* ability to respond to different chemical agents, preparations containing 10^4^, 10^3^, 10^2,^ and 10 freshly harvested spores were spotted on solid YNB media, pH 3.0, supplemented with SDS (0.002%), and EMS (0.05%). In addition, 200 fresh spores plated on YNB agar plates with pH 3.0 were irradiated with UV (10 mJ/cm^2^) to evaluate the fungal survival ability. All fungal cultures were incubated at 26°C, and after 48 h, the survival percentage was calculated from the total number of cells obtained in control conditions.

### *In vitro* host-pathogen interaction assays

For the host-pathogen interaction experiments, the fungal spores were challenged with mouse macrophages (J774A.1) according to previously established protocols [68]. In brief, the freshly harvested spores of the *set1* mutants were added to cell cultures, in a proportion of 1.5 spores per macrophage, in Leibovitz L-15 media (Biowest, Minneapolis, MN, USA) amended with fetal bovine serum bovine (10%) and 1% penicillin/streptomycin (Gibco) and incubated at 37°C. After 30 minutes of co-incubation, non-phagocytized spores were removed, adding phosphate-buffered saline (PBS, 1X) to the cultures. After 5.5 h of interaction, pictures of germinated spores in the macrophages were taken to determine the polarity index using ImageJ software, as reported previously [68]. To evaluate the *M. lusitanicus* survival, the phagocytized spores were released by the addition of NP-40 (0.1%, Sigma Aldrich) to lyse cells. The recovered spores (500 spores) were plated on MMC plates with pH 3.2, placed at 26°C, and after 48 h, the development of healthy colonies was visually inspected. Spores not confronted with macrophages and macrophage cultures served as control samples.

### Determination of cell death

For the cell death experiments, spore-macrophage interactions were prepared in the same conditions mentioned above [68]. After 24 h of co-incubation at 37°C, propidium iodide (PI, Invitrogen) was added to the co-cultures to evaluate macrophage death, according to supplier recommendations. The images were taken with Texas Red and bright-field filters 30 minutes after PI application using the 20/0.8-A objective from a Nikon Eclipse 80i fluorescent microscope equipped with a Nikon DS-Ri2 camera. The images were processed into binary images and overlapped using ImageJ software. The cell death ratio was estimated by manually counting the PI-stained cells and the total number of macrophages in the microscope images.

### Infection assays in *Galleria mellonella*

Sixth instar larvae of *G. mellonella* (SAGIP, Italy), weighing 0.3-0.4g, were selected for experimental use [81]. 10^6^ spores in a volume of 20 µl were injected per larva through one of the hind pro-legs as described previously [82]. Larvae were incubated at 30°C. Untouched larvae and larvae injected with sterile IPS (insect physiological saline [83]) served as controls. *G. mellonella* survival was monitored every 24 h up to 144 h. For each test group, 20 larvae were used, and experiments were repeated at least twice. Survival curves were statistically analyzed by log-rank (Mantel-Cox) test, utilizing GraphPad Prism 8.0.2 software. P values ≤ 0.05 were considered statistically significant.

### RNA-sequencing analysis

Total RNA was extracted from MU636 and *set1* mutant strains growing in plates using the RNeasy Plant Mini Kit (Qiagen) and treated with DNase (Sigma, On-Column DNaseI treatment set). RNA integrity was quality checked using the Bioanalyzer 2100 (Agilent), and samples were submitted to Novogene for library preparation and sequencing. Raw paired-end (PE) reads from *M. lusitanicus* RNA-seq datasets were quality-checked using FastQC v0.11.9 before and after removing adapter sequences with Trim Galore! V0.6.7. Pairs containing a read with a Phred quality score (q) ≤ 33 and/or a total length < 20 nt were removed from the analysis, as well as adapter sequences with an overlap ≥ 4 nt. The PE processed reads were aligned to *M. lusitanicus* v3.0 genome (herein Mucci3, available at https://mycocosm.jgi.doe.gov/Mucci3/Mucci3.home.html) using hisat2 v2.2.1 [84] with a maximum intron length of 500 nt. Individual count matrices were created from the Binary Alignment Maps (BAM) using featureCounts v2.0.3 [85], excluding multimapping reads (Supplementary data ‘Raw counts’). PE reads were specified (p) and counted as fragments (countReadPairs) at the gene level (t) with protein ID as attribute (g). Differential gene expression analysis was performed by DESeq2 v1.34.0 package [86]. Genes with a False Discovery rate (FDR) ≤ 0.05 and a log2 fold change (log2FC) ≤ 1 or ≥ 1 were considered differentially expressed genes (DEGs). The volcano plot of DEGs was generated by VolcaNoseR web app [87].

### Protein prediction and structural analysis

Structure prediction of the MlSac1 protein (ID1537005) was performed using AlphaFold2 [88] implemented with the freely available ColaFold pipeline [89]. ColabFold was run with the default configuration, except for selecting template mode with pdb70 database, and the multiple sequence alignments were generated with MMseqs2 [90]. MlSac1 model 1 was superimposed to ScSac1 (PDB <u>3LWT</u>) by Matchmaker using ChimeraX [91].

## Data availability

The raw data and processed files generated in this work are deposited at the Gene Expression Omnibus (GEO) repository and are publicly available through the project accession number GSE233894.

## Author contributions

Conceptualization: V.G and M.O.C.

Formal analysis, M.O.C., C.L., U.B., G.T., J.T.C.-M., and D.L.G.

Funding acquisition, V.G. and F.E.N. Investigation, M.O.C., C.L., G.T., E.N., and D.L.G. Resources: U.B. and E.N.

Supervision: V.G.

Validation: U.B., F.E.N., V.G.

Writing – Original Draft Preparation: M.O.C., F.E.N., and V.G.

Writing – Review & Editing: M.O.C., C.L., U.B., G.T., J.T.C.-M., D.L.G., E.N., F.E.N., and V.G.

All authors have read and agreed to the published version of the manuscript.

**Conflicts of Interest:** The authors declare no conflict of interest. The funders had no role in the design of the study; in the collection, analyses, or interpretation of data; in the writing of the manuscript, or in the decision to publish the results.

## Figure captions

## Supporting information

**S1 Table. Set1 proteins used for phylogenetic analysis.**

**S2 Table. Mutants of *M. lusitanicus* generated in this study.**

**S3 Table. Primers used in this study.**

**S1 Fig. *set1* knockout strains of *M. lusitanicus* are sensitive to SDS and EMS.**

**S2 Fig. Phenotypic analysis of *set1* knockout strains of *M. lusitanicus* during the interaction with mouse macrophages (J774A.1).**

**S1 Dataset. Proteins and sequences used in the conservation analyses.**

**S2 Dataset. DEGs in *set1* mutants**

**S1 Fig.**
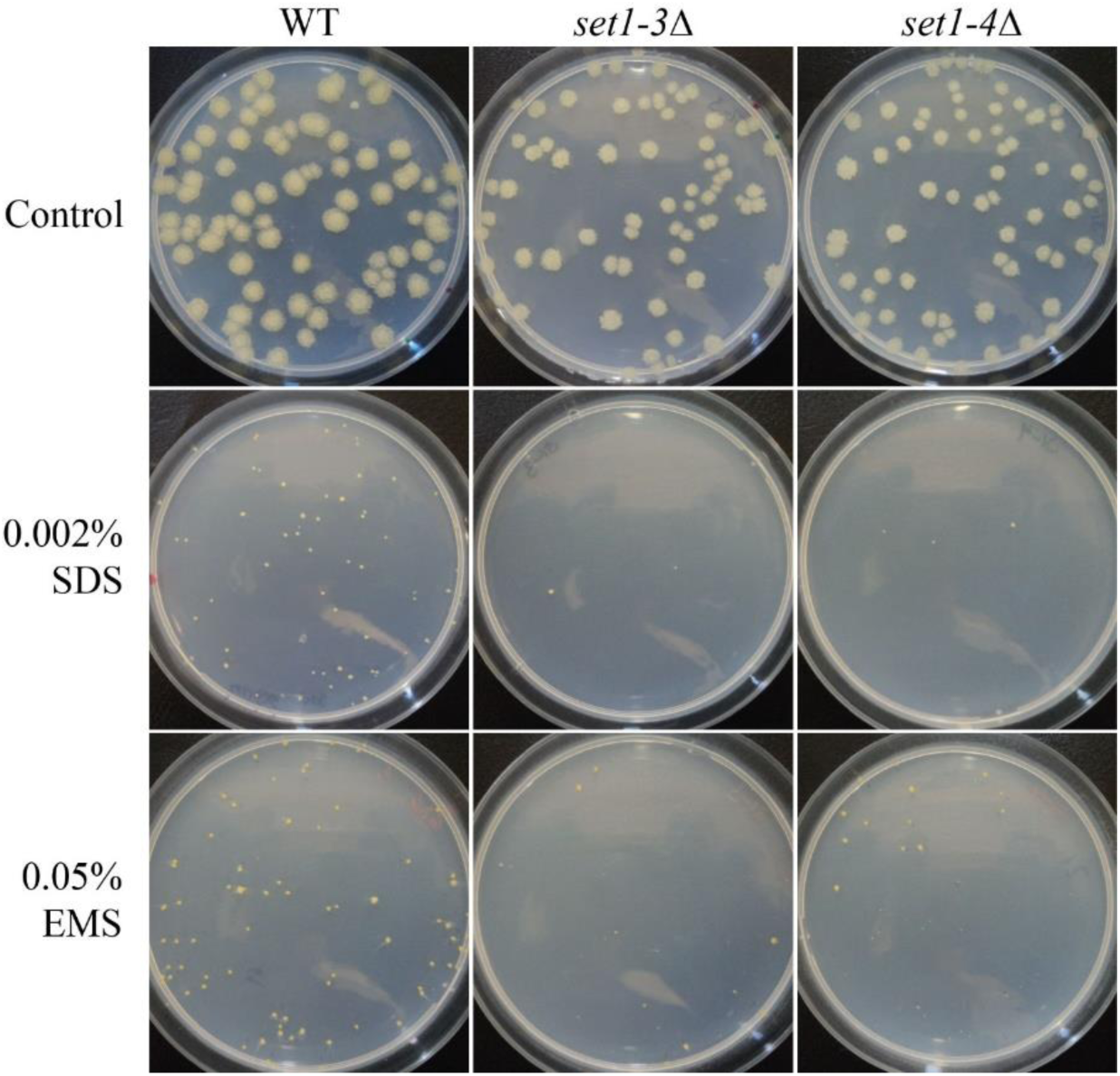
***set1* knockout strains of *M. lusitanicus* are sensitive to SDS and EMS**. Fungal Colonies of MU636 (WT), *set1-3*Δ and *set1-4*Δ with 48 hours of growth on YNB plates (Control) and YNB medium supplemented with SDS (0.002%) or EMS (0.05%) at 26°C.

**S2 Fig.**
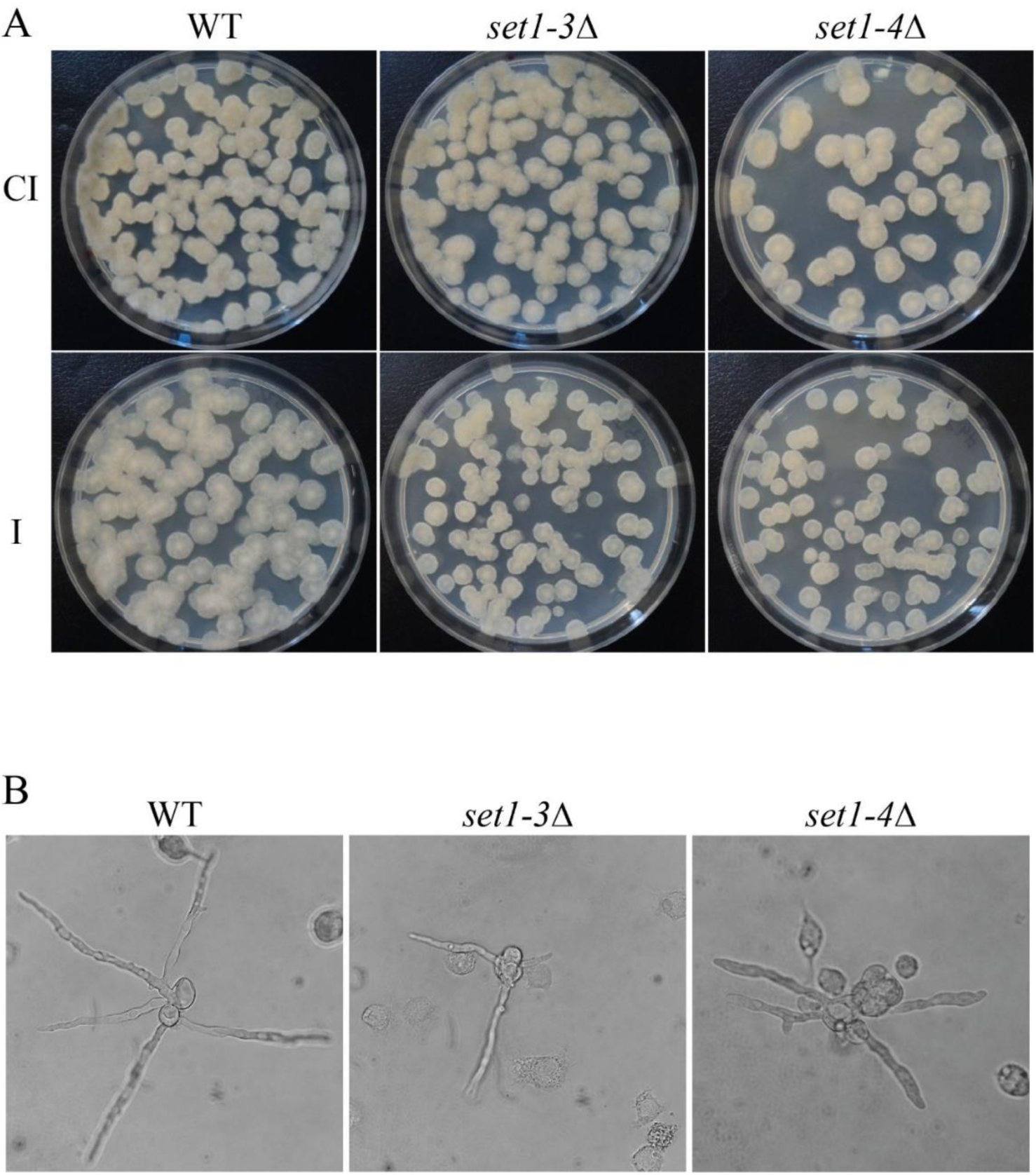
Phenotypic analysis of *set1* knockout strains of *M. lusitanicus* during the interaction with mouse macrophages (J774A.1). A) Colonies of wild-type strain MU636 (WT) and *set1* deletion mutants developed from spores undergo phagocytosis by mouse macrophages during 5.5 hours of co-culture at 37°C (I). The released spores from macrophages were plated on MMC plates to check the growth and colony morphology at 48 hours of culture at 26° C. Spores cultivated under the same conditions but without macrophages served as controls (CI). B) Images show the emergence of the germ tube from spores of WT and *set1*Δ strains phagocytosed by mouse macrophages after 5.5 h of co-cultivation.

**Table S1.** Set1 proteins used for phylogenetic analysis.

| **Name** | **Specie** | **ID FungiDB** | **ID JGI** | **ID NCBI** |
| --- | --- | --- | --- | --- |
| Nc | *Neurospora crassa* OR74A | NCU01206 |  |  |
| Fg | *Fusarium graminearum* PH-1 | FGRAMPH1_01G24837 |  |  |
| An | *Aspergillus nidulans* FGSC A4 | AN5795 |  |  |
| Ca | *Candida albicans* SC5314 | C1_00960C_A |  |  |
| Sc | *Saccharomyces cerevisiae* S288C | YHR119W |  |  |
| Ml | *Mucor lusitanicus* CBS 277.49 |  | 137855 |  |
| Rm | *Rhizopus microspores* ATCC11559 |  | 286187 |  |
| Ao | *Apophysomyces ossiformis* |  |  | KAF7725527 |
| Ar | *Amylomyces rouxii* NRRL 5866 |  | 625830 |  |
| At | *Arabidopsis thaliana* |  |  | OAP08766 |

**S2 Table.** *M. lusitanicus* strains used in this study.

| **Name** | **Genotype** | **Origin** |
| --- | --- | --- |
| MU402 | *pyrG<sup>-</sup>, leuA<sup>-</sup>* | [1] |
| MU636 | *leuA<sup>-</sup>* | [2] |
| *set1-3Δ* (MU1350) | *set1Δ::pyrG<sup>+</sup>, leuA<sup>-</sup>* | This work |
| *set1-4Δ* (MU1350) | *set1Δ::pyrG<sup>+</sup>, leuA<sup>-</sup>* | This work |

---

**S3 Table.**
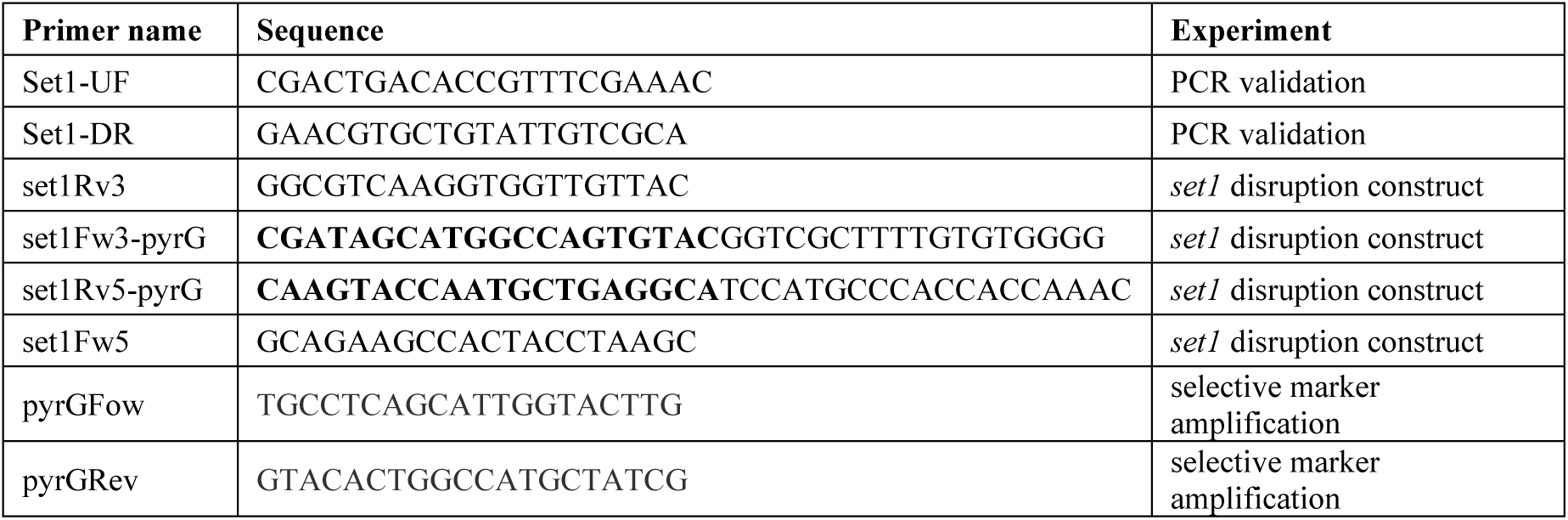
Primers used in this study

